# Peptidylglycine α-amidating monooxygenase restores brain microvascular blood flow and improves recovery following ischemic stroke

**DOI:** 10.64898/2026.06.14.732201

**Authors:** Ella Matson, Yulia Illina, Anania Tsinoglou, Emily H Attrill, Sophie Mayne, Renee M Ross, Michelle A Keske, Brad A Sutherland, Harald Hampel, Paul Kaufmann, Andreas Bergmann, Dino Premilovac

## Abstract

**Introduction:** Reduced or absent capillary blood flow (termed no-reflow) even after arterial recanalization is associated with poorer neurological outcomes following ischemic stroke. The aim of the current study was to test whether acute administration of peptidylglycine α-amidating monooxygenase (PAM) can increase capillary blood flow and improve brain recovery after ischemic stroke.

**Methods:** A 60-minute ischemic stroke was induced using middle cerebral artery occlusion (MCAO) in rats. A modified, long-acting PAM enzyme was administered 30 minutes after induction of ischemia and rats were recovered for either 24 hours or 7 days. In all animals, real-time cerebral blood flow was assessed before, during and after MCAO using trasncranial contrast enhanced ultrasound (tCEU). For rats in the 7-day protocol, a modified neuroscore test was used to assess neurological deficit following MCAO. At the end of each experiment, a transcardiac perfusion was used to generate a fluorescent vascular cast and histology was used to examine capillary diameters and determine infarct volume.

**Results:** Following MCAO and arterial recanalization, untreated rats had reduced cerebral blood flow across brain regions affected by ischemia, indicative of no-reflow. PAM administration led to enhanced cerebral blood flow in affected regions, and this was associated with increased capillary diameters 24 hours after ischemic stroke. Although there was no difference in infarct volume at 24 hours, by day 7, infarct volume was markedly reduced in the PAM group and these animals exhibited improved neurological function compared to the untreated group.

**Conclusion:** Administration of PAM improves capillary blood flow after ischemic stroke leading to enhanced neurological and brain recovery. This work highlights PAM as a novel theraputic approach to improve brain blood flow and recovery after ischemic stroke.

**GRAPHICAL ABSTRACT:** 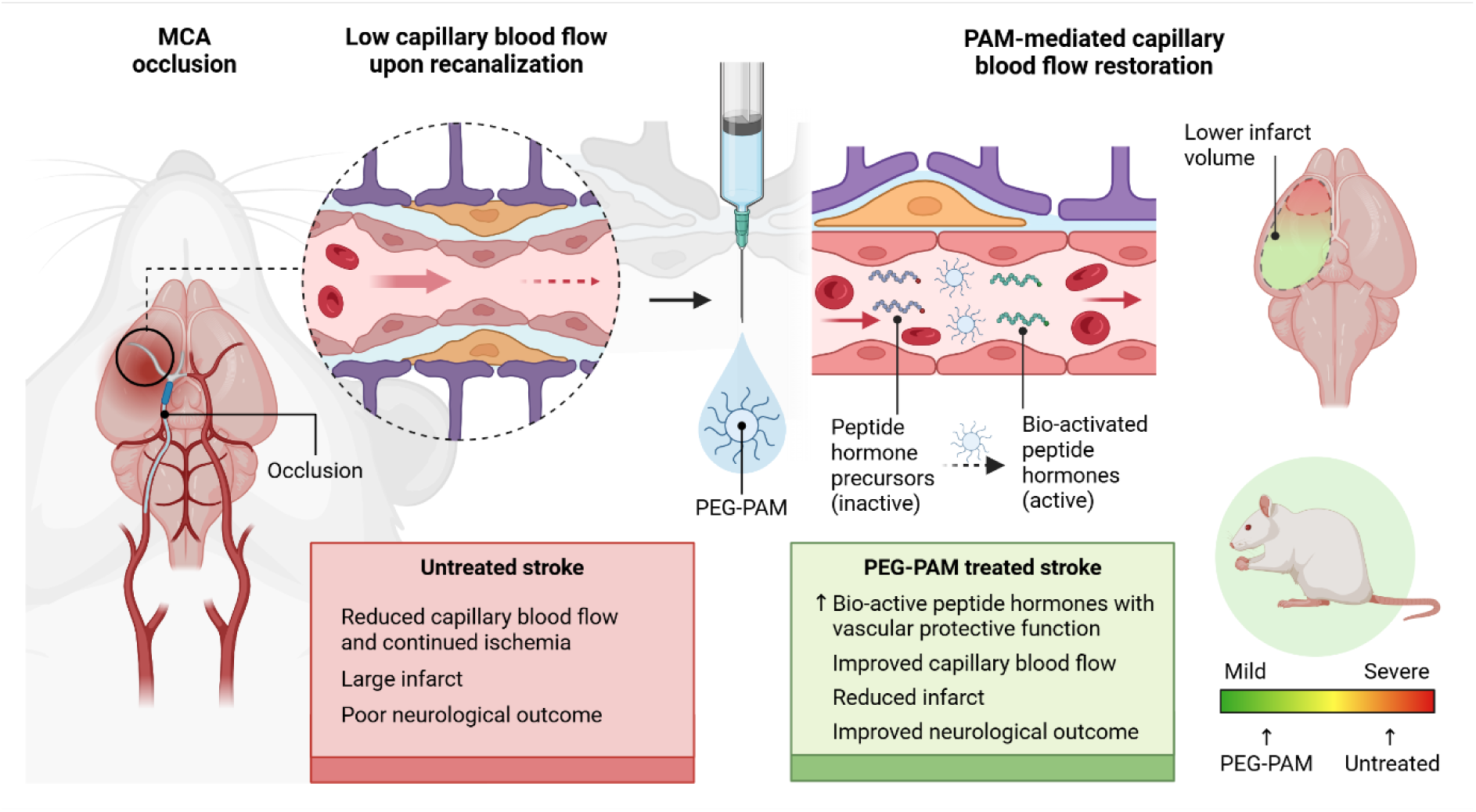

## INTRODUCTION

Current acute ischemic stroke therapies, intravenous thrombolysis (alteplase) or endovascular thrombectomy, work to restore large-vessel patency and improve outcome (1, 2). However, up to 50% of patients still have poor functional recovery despite successful arterial recanalization (1, 2). Preclinical and clinical studies implicate microvascular failure as a key reason, where obstruction and dysregulation at the capillary level limits tissue perfusion even after the arterial occlusion is removed (termed no-reflow or the futile reperfusion phenomenon) (3, 4). Capillary no-reflow is attributed to mechanisms such as leukocyte or microthrombus plugging, pericyte-mediated capillary constriction, and endothelial barrier disruption (4–6). Additionally, inflammation and oxidative stress after reperfusion disrupt the blood brain barrier (BBB) and promote vasogenic edema, further narrowing capillaries and contributing to capillary no-reflow and continued ischemia (7, 8).

The no-reflow phenomenon is common in both humans and animal models of ischemic stroke (5, 9–11). Clinical studies report a ∼20-50% incidence of focal hypoperfusion after successful recanalization (9–11). This sustained reduction in capillary blood flow in damaged brain regions drives continued ischemia and is associated with poor neurological recovery and increased risk of hemorrhagic transformation (9, 11). Therefore, re-establishing normal capillary blood flow acutely after ischemic stroke treatment is critical to improve subsequent brain recovery and patient outcomes.

One potential avenue to support microvascular recovery after stroke is to enhance the levels of endogenous vasoactive and neuroprotective peptides, many of which must be enzymatically amidated to become bioactive. Peptidylglycine α-amidating monooxygenase (PAM) is the only known enzyme capable of catalyzing C-terminal α-amidation, a post-translational modification required for the biological activity of more than 70 peptide hormones (12). PAM substrates include adrenomedullin (ADM), calcitonin gene-related peptide (CGRP), vasoactive intestinal peptide (VIP), pituitary adenylate cyclase-activating polypeptide (PACAP) and neuropeptide Y, many of which exert vasodilatory and neuroprotective effects relevant to cerebral ischemia (12, 13).

Several amidated peptide hormones are dysregulated following ischemic stroke and appear to form part of the endogenous response to vascular injury (14, 15). ADM, CGRP, VIP, PACAP and related peptides have been implicated in the regulation of cerebral blood flow, endothelial function, blood-brain barrier integrity and neuroinflammation (16–18). Experimental administration of several of these peptides improves cerebral perfusion, BBB stability and neurological recovery in preclinical stroke models, suggesting that amidated peptide signalling contributes to endogenous recovery mechanisms following cerebral ischemia (19–21). Despite this biological rationale, the therapeutic use of individual peptide hormones remains challenging. Most peptide hormones exhibit rapid plasma clearance, and require continuous or repeated intravenous administration to maintain therapeutic concentrations (22, 23). An alternative approach is therefore to enhance endogenous peptide activation rather than supplementing individual hormones. PAM occupies a unique position in this regard, as it is the sole enzyme responsible for C-terminal α-amidation. Augmenting PAM activity directly in the circulation could simultaneously increase the activation of multiple vasoactive peptide systems while preserving their physiological regulation in the context of ischemia.

Although peptide amidation has historically been considered a process restricted to secretory granules (13, 24), recombinant PAM has been shown to retain catalytic activity in the circulation and convert glycine-extended peptide precursors into their bioactive amidated forms *in vivo* (25). Furthermore, substantial circulating pools of glycine-extended substrates have been demonstrated for several PAM-dependent hormones, including ADM and gastrin, indicating that the bloodstream contains both the substrate and the necessary cofactors required for ongoing amidation (25, 26). These findings suggest that the endogenous circulation itself can serve as a therapeutically accessible compartment for PAM-mediated peptide activation.

Previous work demonstrates that recombinant human PAM remains enzymatically active in the circulation and increases systemic amidating activity *in vivo* (27). However, native PAM has a short plasma half-life (42–47 minutes), limiting its therapeutic applicability (27, 28). PEGylation of PAM (PEG-PAM) extends its half-life to approximately 218 minutes and enables sustained *in vivo* amidating activity for up to 7 days following a single administration (28). Sustained augmentation of PAM activity therefore represents a strategy to support endogenous activation of multiple, ischemic-damage relevant vasoactive peptides without requiring continuous peptide infusion. This study is the first to investigate the theraputic potential of sustained PAM activity, via a modified enzyme, in the setting of ischemic stroke. We hypothesize that PEG-PAM administration improves capillary perfusion by enhancing circulating amidating capacity during the post-ischemic period and that this will lead to improved brain recovery. The primary aim of this study was to determine whether PEG-PAM administration improves microvascular flow and brain recovery following experimental ischemic stroke.

## METHODS

### Animal Husbandry and ethical approval

Male Sprague Dawley rats at 10-12 weeks old were used for all experiments. All rats were housed in standard conditions (21±2°C, 12hr-12hr light-dark cycle) and were provided water and chow *ad libitum*. All rats were randomly allocated to one of the following groups; i) sham surgery, ii) middle cerebral artery occlusion (MCAO) plus vehicle (MCAO+Vehicle), or, iii) MCAO plus PEG-PAM (MCAO+PAM). Animals in these three groups were then allocated to different experiments to understand the effect of PEG-PAM administration on short term recovery (24 hours; experiment 1) or longer-term recovery (7 days; experiment 2) following ischemic stroke – see Figure 1. In both experiments, vehicle or PEG-PAM (200µg/kg) were administered 30 minutes prior to arterial recanalization. All experimental procedures were approved by the University of Tasmania Ethics Committee (A0028593) and performed in accordance with the Australian Code for the Care and Use of Animals for Scientific Purposes – 2013 (updated 2021), 8^th^ Edition.

**Figure 1.**
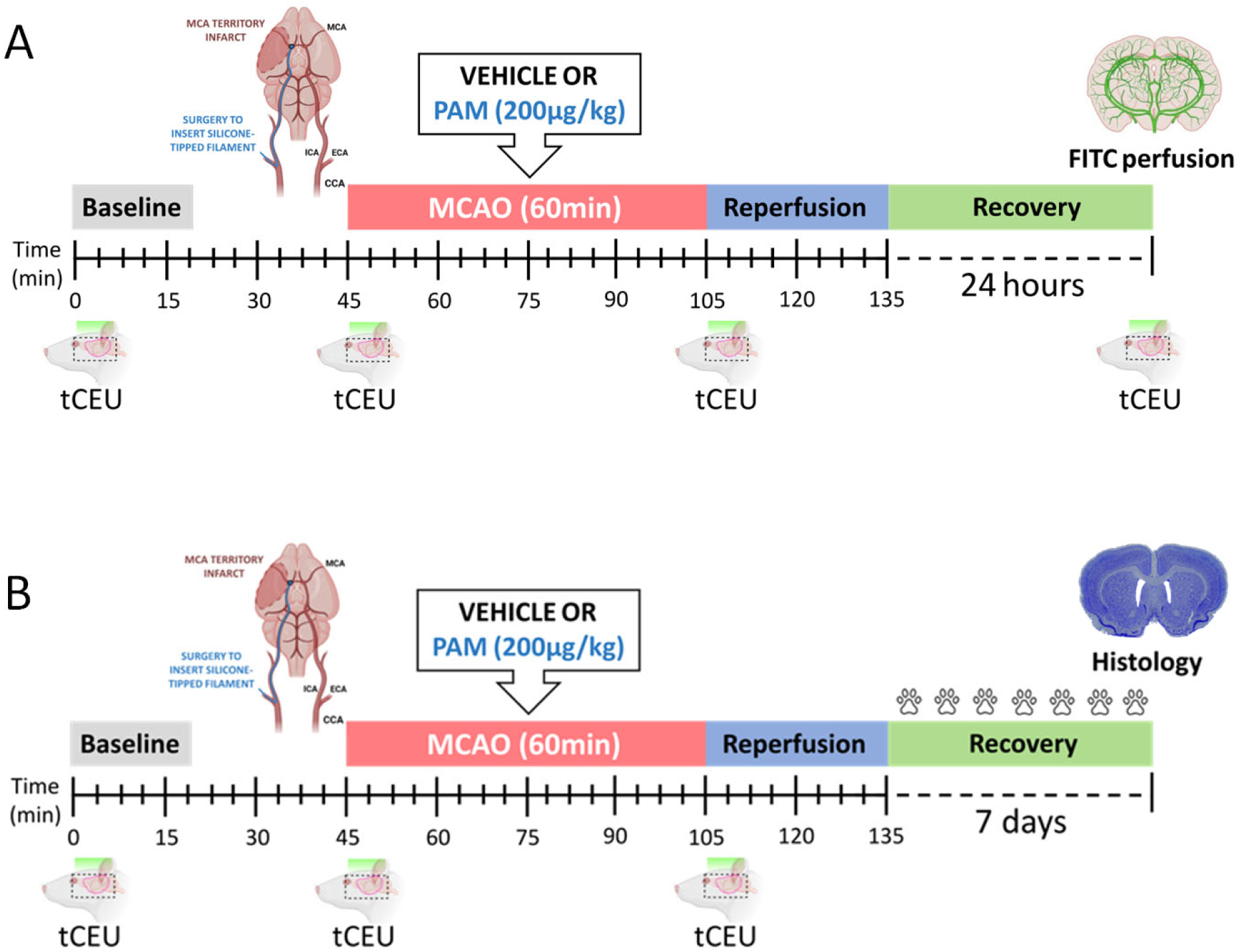
Experimental timeline for the 24-hour recovery and 7-day recovery experiments. Panel. **A.** For the 24-hour recovery experiment, all rats were anesthetized with isoflurane. The carotid artery was cannulated with a silicone tipped filament to induce middle cerebral artery occlusion (MCAO) to mimic ischemic stroke. The filament was kept in place to induce ischemia for 60 minutes. After 30 minutes of ischemia, animals were administered an intraperitoneal injection of vehicle (saline) or PEG-PAM (200µg/kg). After 60 minutes of ischemia, the filament was withdrawn to enable arterial recanalization and tissue perfusion. tCEU imaging was used to measure blood flow at baseline, MCAO, immediately after reperfusion. After this, animals recovered for 24 hours after which one final tCEU measurement was performed and animals underwent transcardiac perfusion with 4% PFA followed by gelatin containing FITC-dextran to create an *in situ* fluorescent vascular cast. **Panel B.** For the 7-day recovery experiment, all rats underwent the same procedure as in the 24-hour experiment but were recovered for 7 days following the 60-minute MCAO. As in experiment 1, rats were treated with vehicle (saline) or PEG-PAM (200ug/kg) 30 minutes prior to filament withdrawal and restoration of arterial blood flow. tCEU was used to measure blood flow changes at baseline, during MCAO and at acute reperfusion. After recovery, a modified neuroscore was used to assess motor and sensory function each day for 7 days after MCAO. At the end of the 7 days, brains were collected and infarct volume determined using cresyl violet staining.

### PAM Constructs and PEGylation

The recombinant human PAM construct used in this study comprises the soluble full-length sequence (Met1-Ser866) lacking the protease-sensitive linker region (aa 388–494). The construct was expressed in CHO-S cells and includes a C-terminal His-tag. Recombinant PAM was purified using nickel-affinity chromatography under standard conditions. PEGylation of soluble PAM was performed using N-hydroxysuccinimide-activated methoxy polyethylene glycol (5 kDa) according to a previously described protocol (28). The resulting PEG-PAM retained amidating activity and was used for all *in vivo* experiments. Circulating PEG-PAM concentrations were quantified in terminal plasma using a validated sandwich immunoassay, as previously described (29).

### Vascular surgery and ischemia induction

MCAO surgery was performed as previously published (30). Briefly, following successful isoflurane anesthesia, neck microsurgery was performed to cannulate the external branch of the right jugular vein to enable intravenous infusion of phospholipid microbubbles for transcranial contrast enhanced ultrasound (tCEU) blood flow imaging. After this, the right external carotid (ECA), internal carotid (ICA), and common carotid arteries (CCA) were carefully exposed and isolated. The ECA was permanently cauterized and cut to form a stump and the CCA and ICA were temporarily ligated. A small arteriotomy was made in the ECA stump, and a silicon-tipped filament (404156, Doccol Corporation, USA) was inserted into the ECA and carefully advanced up the ICA until resistance was felt, indicating the filament was lodged at the origin of the middle cerebral artery (MCA). After 60 minutes of ischemia, the filament was withdrawn and the ECA cauterized to seal the incision site. Lastly, the CCA ligature was released to allow full recanalization of the arterial system, mimicking endovascular thrombectomy (31). Lastly, the surgical site was sealed using non-absorbable sutures and animals recovered for either 24 hours or 7 days.

### Non-invasive brain blood flow imaging using tCEU

We have previously optimized tCEU to measure real-time blood flow in the brain without requiring a craniotomy or any incisions into the head (30). In brief, an ultrasound transducer (L12-5) was positioned over the right side of the head and interfaced with an iU22 ultrasound machine (Philips Medical Systems, Australia). Real-time (15.2Hz), low mechanical index (0.24) imaging was performed during a constant (55µl/min) infusion of phospholipid microbubbles (Definity®, Lantheus Medical Imaging, Australia, 1ml of activated Definity® added to 5ml saline) at each imaging period. The ultrasound gain (94%), compression (C=39), depth (2 cm) and focus were kept constant for all imaging sessions across all rats.

We used tCEU to quantify brain blood flow before MCAO, during MCAO, 5 minutes after recanalization (acute reperfusion) and 24 hours after recanalization (experiment 1 only). To ensure consistency in imaging at each timepoint, a 10-minute infusion of microbubbles was used to ensure steady-state arterial concentration. After this, a series of high mechanical index (1.06) pulses were used to destroy all the microbubbles in the vasculature under the transducer. The refill of microbubbles over 60 seconds through the vasculature was recorded and this sequence was repeated three times at each point in the protocol. All recordings were analyzed using QLab^TM^ (version 13.0, Philips Medical Systems) imaging software. A rat sagittal brain atlas (32) was superimposed over the sagittal tCEU images to identify the cortical and subcortical brain regions for quantification. These regions are primarily supplied by the MCA and therefore undergo ischemia during MCAO followed by reperfusion after filament withdrawal. For each brain region, the three 60 second recordings were averaged and fitted to an exponential increase to maximum function in Sigma Plot 11 (Systat Software Inc, San Diego, CA, USA) using the following equation:

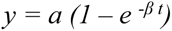

Here, *y* = acoustic intensity (AI), *a* = vascular volume, *β* = flow velocity, and *t* = time in seconds. Vascular perfusion, reflecting changes in cerebral blood flow, was then calculated as the product of *a* x *β* as previously published (30, 33–35).

### Assessment of neurological deficit

To assess neurological function post-MCAO in the 7 day recovery experiment, all rats underwent daily neurological scoring to determine level of motor and sensory deficit (36, 37). The neuroscore assessment used in this study is based on the modified Rankin Scale, which is the most commonly used outcome measure in clinical stroke trials (38). Neuroscore consisted of an assessment of the rats’ general activity, symmetry of movement, forelimb preference, whisker stimulation response, trunk touch response, and climbing ability, all ranked on a scale of 0 (no impairment) to 3 (severe impairment). The scores for each category were then summed to give a score out of 18, with a higher neuroscore corresponding to a higher level of motor and/or sensory impairment. Assessors were blinded to interventions where possible over the 7-day assessment period.

### Brain histology and infarct analysis

After the end of each experiment, all rats were injected with a terminal dose of sodium pentobarbitone (>200mg/kg). After deep anesthesia was confirmed, the heart was exposed and an 18G needle was inserted into the left ventricle to facilitate transcardial perfusion with heparinized phosphate buffered saline (PBS) via a peristaltic pump to clear blood from vasculature. After this, transcardial perfusion switched to 4% paraformaldehyde (PFA; pH 7.4) in PBS for 5 minutes followed by fluorescein isothiocyanate conjugate (FITC)-albumin gelatin solution (1.25% w/v gelatin, 0.1% FITC-albumin in PBS; Merck, USA, #A771-1G). The flow rate for transcardial perfusion was adjusted for each animal to maintain a systemic perfusion pressure of 80-120mmHg (measured using an in-line sphygmomanometer) to mimic physiological perfusion pressure *in vivo*. Once perfusion ceased, the rats were covered in ice for 30 minutes to allow the gelatin to set, creating an intravascular fluorescent cast as described (39). After this, brains were carefully excised and cryoprotected using 30% sucrose in PBS at 4°C for three days. The brains were then set in optimal cutting temperature compound (OCT; Agilent Technologies, USA). All brains were serially sectioned using a cryostat at -20°C to generate a series of 40µm sections for each brain. To quantify the extent of damage caused by ischemia, infarct volume was measured using cresyl violet acetate. To do this, a series of sections from each brain (1 section per 1mm; 9 sections total per brain) was stained in a 0.1% cresyl violet acetate solution for 12 minutes and cover-slipped using DPX Neutral Mounting Medium (ChemSupply, SA, Australia, #DL028). Slides were digitized using brightfield imaging on a VS200 Slide Scanner (Olympus, Australia) at 20x magnification. QuPath (version 0.5.1; (40)) was used to annotate total area (µm^2^) of the ipsilateral and contralateral brain hemispheres across each of the serial brain sections to enable quantification of infarct volume. For experiment 1, the Reglodi method (41) was used to determine edema-adjusted infarct volume (lesion area x (contralateral area/ipsilateral area)). After this, a coronal brain atlas was used to identify cortical and subcortical brain regions in each section to enable cortical and subcortical infarct volume quantification.

### Microvascular analysis

For immunohistochemical analysis of brain microvasculature, a series of brain sections from each animal was incubated overnight at 4°C in PBS containing 4% donkey serum, 0.4% triton-X-100 and anti-laminin antibody (1:1000; Abcam, UK; ab11575) to detect blood vessels. After this, sections were washed using PBS and incubated in the dark for 1 hour at room temperature in PBS containing 4’,6-diamidino-2-phenylindole (DAPI; 1:20,000; Merck, USA) and donkey anti-rabbit secondary antibody (Alexa Fluor® 647; 1:1000; Abcam, UK; ab150075). Sections were washed with PBS and glass coverslips were mounted using fluorescence mounting medium (Dako, Denmark). Slides were imaged at 20x magnification using extended focal imaging on an Olympus VS200 Virtual Slide System (DAPI:388nm, 50ms; FITC-albumin:488nm, 100ms; laminin: 647nm, 200ms; Olympus, Japan) and analyzed using ImageJ (version 1.52i; NIH, USA) and QuPath.

To understand how MCAO with and without PEG-PAM treatment impact capillary no-reflow, the intraluminal FITC signal was quantified to estimate total perfused area as well as direct measurement of capillary lumen diameter. For estimation of total perfused area, 3000×3000 pixel (1035×1035µm) regions of interest were extracted from either the cortical or sub cortical brain regions on both the ipsilateral (MCAO) and contralateral brain hemispheres. For each region, the total number of FITC pixels was calculated as a percentage of all pixels in the region of interest. The contralateral brain region was used as the internal control for each animal by comparing total FITC pixels in the corresponding ipsilateral region to the contralateral (expressed as % of the contralateral). In addition to this, 10 capillaries in each region were selected using the laminin signal for manual interrogation of capillary wall and lumen diameters. To avoid bias, the operator was blinded, and selection and analysis was performed in a randomized manner using a random point generator. Manual measurements of capillary wall (laminin; cyan) and capillary lumen (FITC; green) diameter were completed using the line drawing tool in QuPath. Capillary wall diameter was measured at the midpoint of the capillary and capillary lumen diameter was assessed ∼1-2µm on either side of the midpoint with the subsequent measurements averaged into one number per capillary. The mean capillary diameter from each animal for each brain region was calculated from the 10 individual capillaries.

### Data analysis and Statistics

All animals were randomized to treatments and researchers were blinded throughout the experiments where possible for all interventions and measurements. Statistical analysis was performed using Sigma Plot version 11.0 (Systat Software Inc). All data was tested for normal distribution using the Shapiro-Wilk test. When data was not normally distributed, the BOXCOX transformation was used to normalize data before parametric testing (42). Single time-point and immunohistochemical data were assessed using a one-way ANOVA with Student-Newman-Keuls *post hoc* test. When measurements were repeated across time two-way repeated measures ANOVA was used followed by the Student-Newman-Keuls *post hoc* test. For neuroscore data, the Kruskal-Wallis test was used to determine differences between groups at day 7. All data presented as mean and standard deviation and individual data points are shown in graphs where relevant. A p-value <0.05 was considered statistically significant.

## RESULTS

### PEG-PAM administration does not alter systemic hemodynamics or metabolism in healthy rats

First, we assessed the potential effect of PEG-PAM on systemic hemodynamics and markers of metabolism in healthy rats. To do this, rats received intraperitoneal injections of PEG-PAM (200µg/kg) and changes in mean arterial blood pressure and heart rate were recorded throughout and arterial blood samples were collected every 20 minutes (Supplementary Figure 1A). Compared to vehicle administration, PEG-PAM administration did not alter mean arterial blood pressure (Supplementary Figure 1B) or heart rate (Supplementary Figure 1C) at any timepoint. Similarly, PEG-PAM administration did not alter blood glucose (Supplementary Figure 1D) or blood lactate (Supplementary Figure 1E) concentrations at any timepoint compared to vehicle.

### PEG-PAM improves microvascular blood flow at acute reperfusion and 24 hours after MCAO stroke

We investigated the potential of PEG-PAM to improve microvascular blood flow in the brain after MCAO stroke. First, we used microvascular imaging (MVI) to assess spatial changes in vascular perfusion within the ischemic brain hemisphere (Figure 2A). Brain perfusion for the sham group did not appear to change across any of the time-points. As expected, the induction of MCAO caused a visible reduction in brain perfusion relative to baseline, particularly within the cortical and subcortical regions. At the acute reperfusion phase, the MCAO+Vehicle group appeared to have a sustained reduction in blood flow within the cortical and subcortical regions, indicative of no-reflow. Conversely, blood flow appeared to increase above baseline following reperfusion for the MCAO+PAM group. After the 24-hour recovery, brain perfusion appeared to remain lower than baseline for the vehicle-treated group. In contrast, brain perfusion at 24 hours appeared to return to baseline within the PAM-treated group.

**Figure 2.**
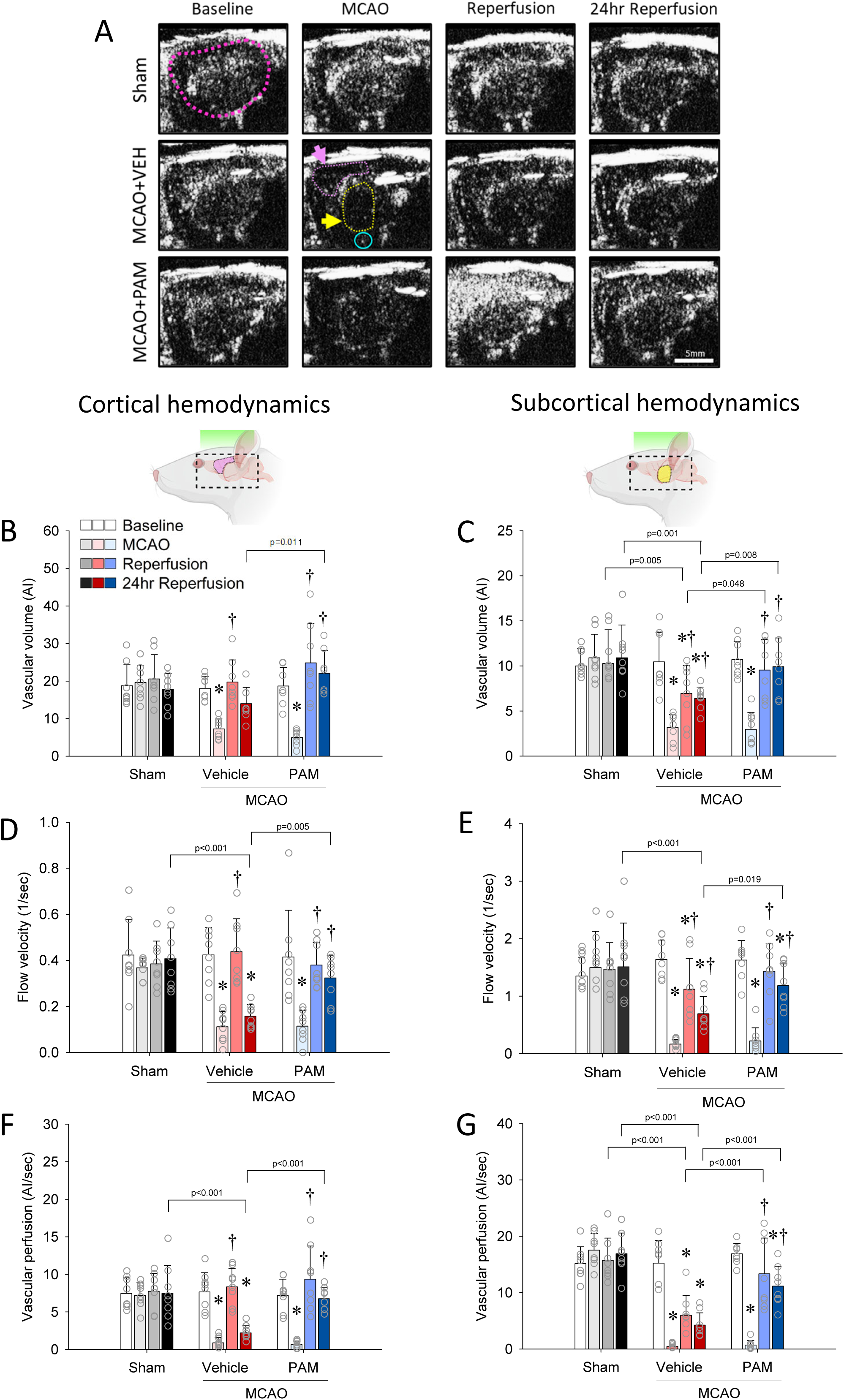
PEG-PAM reduces development of no-reflow in both cortical and subcortical brain regions after 60-minute MCAO. Panel. **A.** Representative tCEU images from one animal at each time point in each group are shown. Pink dotted line in the top-left panel highlights the location of the brain hemisphere in the sagittal orientation. Pink and yellow arrows highlight cortical and subcortical regions, respectively, where MCAO induces marked deficits in blood flow. Cyan circle highlights the MCA at the base of the brain. Scale bar = 5mm. tCEU quantification of cortical vascular volume **(Panel B)**, flow velocity **(Panel D),** and vascular perfusion **(Panel F)** at baseline, MCAO, reperfusion and 24-hour (24hr) reperfusion across all groups. Quantification of subcortical vascular volume **(Panel C)**, flow velocity **(Panel E)**, and vascular perfusion **(Panel G)** at baseline, MCAO, reperfusion and 24hr reperfusion across all groups. Data are means±SD for n=8 animals in each group. Open circles represent data from individual animals. Statistical analysis completed using two-way repeated measures ANOVA with Student-Newman Keuls post hoc. * denotes p<0.05 vs baseline within the group. † denotes p<0.05 vs MCAO within the group. Specific p values are provided for between group comparisons in all relevant panels.

Since the induction of MCAO causes a marked blood flow deficit within the cortical and subcortical regions, we quantified changes in blood flow kinetics specifically within these regions. As expected, the sham group showed no changes in cortical vascular volume, flow velocity, or vascular perfusion across any time-point (Figure 2 B, D and F). The induction of MCAO caused a substantial decrease in vascular volume within the cortex for both the vehicle-treated and PAM-treated groups (Figure 2B; p<0.001 for both vs baseline). Cortical vascular volume then returned to baseline at reperfusion and after 24 hours for both the MCAO+Vehicle and MCAO+PAM groups. Cortical vascular volume at 24 hours was higher in the PEG-PAM-treated group than the vehicle-treated MCAO group (p=0.011).

Flow velocity within the cortex was considerably lower than baseline during MCAO for both the MCAO+Vehicle and MCAO+PAM groups (Figure 2D; p<0.001 for both groups). At reperfusion, cortical flow velocity was restored to baseline levels for both the vehicle-treated and PAM-PAM-treated groups. After 24 hours, we observed a substantial drop in flow velocity within the cortex for the MCAO+Vehicle group when compared to baseline (p<0.001). In contrast, cortical flow velocity for the MCAO+PAM group remained at baseline levels after 24 hours (p=0.196) and higher than the vehicle-treated group (p=0.005).

During MCAO, vascular perfusion within the cortex decreased below baseline, before returning to baseline levels at reperfusion for both the vehicle-treated and PAM-treated groups (Figure 2F; p<0.001 for both groups). After 24 hours, cortical vascular perfusion fell below baseline levels for the MCAO+Vehicle group (p<0.001) but remained at baseline levels for the MCAO+PAM group (p=0.682). At this 24-hour time-point, cortical vascular perfusion was higher in the MCAO+PAM group than the MCAO+Vehicle group (p<0.001).

Next, we analyzed changes in blood flow kinetics within the subcortical region (Figure 2 panels C, E and G). As expected, the sham group showed no changes in subcortical vascular volume, flow velocity, or vascular perfusion across any time-point. The induction of MCAO caused a large reduction in vascular volume within the subcortex for both the MCAO+Vehicle and MCAO+PAM groups (Figure 2C; p<0.001 for both groups). Subcortical vascular volume for the vehicle-treated group failed to return to baseline levels after reperfusion (p=0.002) and at 24 hours (p=0.001). In contrast, vascular volume within the subcortex returned to baseline for the PEG-PAM-treated group at reperfusion (p=0.537) and at 24 hours (p=0.469) and was higher than the MCAO+Vehicle group (p=0.008).

Subcortical flow velocity followed a similar trend to that of subcortical vascular volume (Figure 2E). At the induction of MCAO, there was a substantial drop in flow velocity within the subcortex for both the vehicle-treated group (p<0.001) and PEG-PAM-treated group (p<0.001). Subcortical flow velocity did not return to baseline for the MCAO+Vehicle group at reperfusion (p<0.001) or after 24 hours (p=0.005). In contrast, flow velocity within the subcortex returned to baseline following reperfusion for the MCAO+PAM group (p=0.275). Whilst subcortical flow velocity fell below baseline after 24 hours for the PEG-PAM-treated group (p=0.037), flow velocity remained higher compared with the vehicle-treated group at the 24-hour time-point (p=0.019).

Finally, there was a large decrease in subcortical vascular perfusion for both the MCAO+Vehicle and MCAO+PAM groups during MCAO (Figure 2G; p<0.001 for both groups). Subcortical vascular perfusion remained below baseline for the MCAO+Vehicle group at reperfusion (p<0.001) and after 24 hours (p<0.001), indicative of sustained no-reflow. In contrast, subcortical vascular perfusion returned to baseline levels following reperfusion for the PEG-PAM-treated group (p=0.149). Even though vascular perfusion within the subcortex dropped below baseline after 24 hours for the MCAO+PAM group (p<0.001), subcortical vascular perfusion remained higher than the MCAO+Vehicle group at both reperfusion (p<0.001) and at 24 hours (p<0.001).

### PEG-PAM increases total vascular lumen volume in the cortex but does not modify infarct volume 24 hours after MCAO

Next, we assessed the effect of PEG-PAM on total vascular lumen perfusion within the cortical and subcortical regions. Total vascular lumen volume was quantified using the intraluminal FITC signal by comparing FITC pixels on the ipsilateral and contralateral sides of each brain hemisphere (Figure 3A). In line with the tCEU data, we found that total vascular lumen volume within the cortex was lower for both the MCAO+Vehicle group (p<0.001) and the MCAO+PAM group (p<0.001) when compared to the sham group (Figure 3B). Furthermore, total vascular lumen volume within the cortex was considerably higher in the MCAO+PAM group compared to the MCAO+Vehicle group (p=0.015). Total striatal vascular lumen volume was lower for both the MCAO+PAM (p<0.001) and MCAO+Vehicle (p<0.001) groups relative to the sham group (Figure 3C). However, total striatal vascular lumen volume did not differ between the MCAO+Vehicle and MCAO+PAM groups. Lastly, we used serial brain sections to quantify infarct volume at 24 hours using cresyl violet staining (Figure 3D). As expected, there were no identifiable infarcts within the sham group (Figure 3E). After the 24-hour recovery, total lesion volume did not differ between the MCAO+Vehicle and MCAO+PAM groups. Likewise, there was no difference in lesion volume between MCAO groups when looking specifically within the cortical (Figure 3F) and subcortical (Figure 3G) regions.

**Figure 3.**
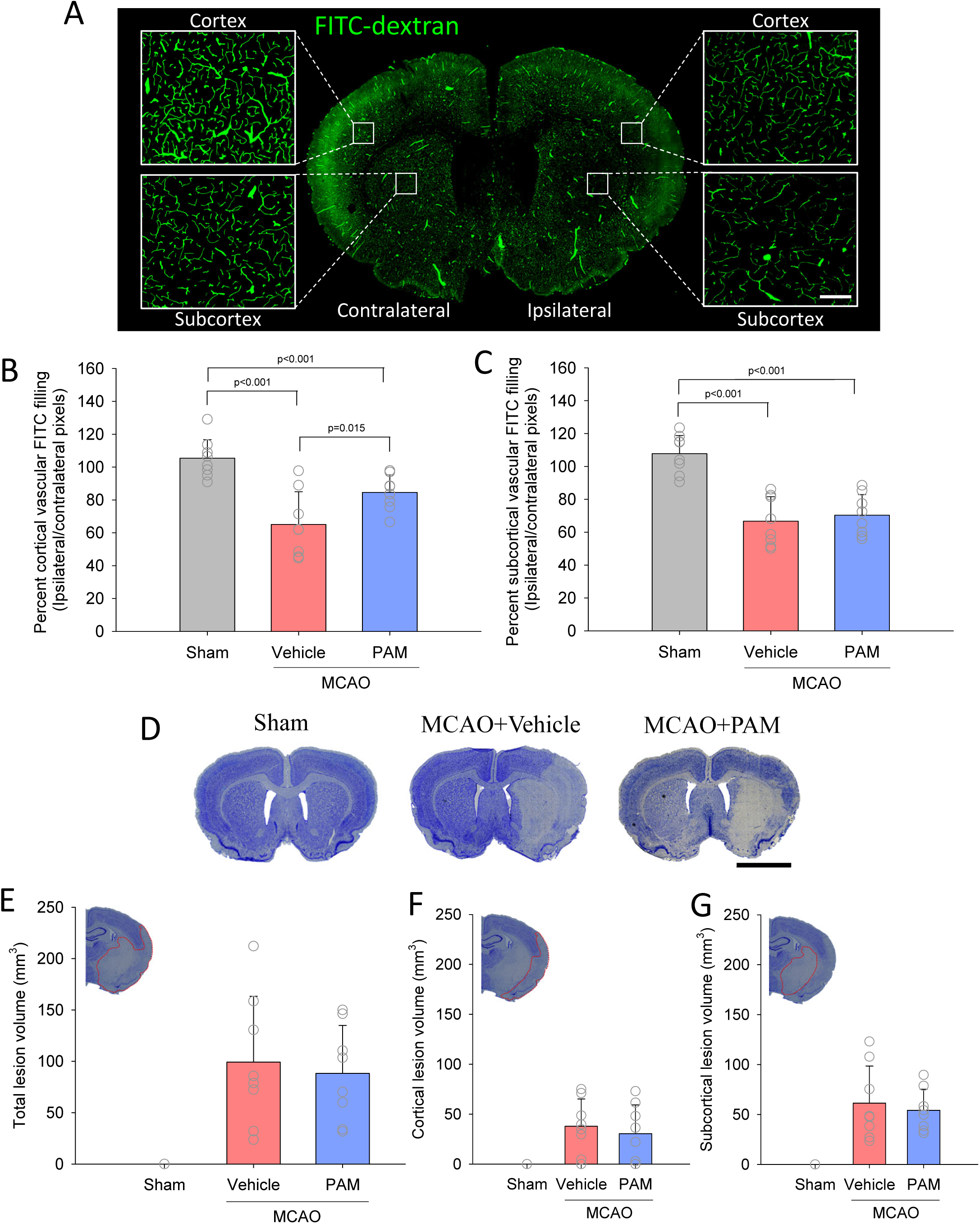
PAM increases total vascular lumen volume in the cortex but does not modify infarct volume 24 hours after MCAO. Panel. **A.** Representative coronal brain section showing FITC-dextran filling of vasculature on the contralateral (no MCAO) and ipsilateral (MCAO) sides. Insets show higher magnification of the cortical and striatal regions on both the contralateral and ipsilateral sides. Scale bar = 100µm. **Panels B and C.** Quantification of FITC filling of vasculature within the cortical (B) and striatal (C) regions was completed by comparing total FITC pixels on the ipsilateral and contralateral sides. **Panel D.** Representative images of coronal brain sections stained with Cresyl Violet from each group. Scale bar = 5mm. **Panels E-G.** Quantification of total (E), cortical (F) and subcortical (G) infarct volumes. Data are means±SD for n=8 animals in each group. Open circles represent data from individual animals. Statistical analysis completed using one-way ANOVA with Student-Newman Keuls post hoc. Specific p values are provided for between group comparisons in all relevant panels.

### PEG-PAM stimulates capillary dilation and reduces capillary no-reflow in the cortical and striatal regions 24 hours after MCAO

To understand whether the vascular improvements noted by tCEU were due to large vessel or capillary changes, we performed a manual assessment of capillary perfusion within the cortical and striatal regions by interrogating capillary wall and lumen diameters (Figures 4 and 5). Overall, capillary wall diameters within the cortex were similar between the contralateral and ipsilateral hemispheres for all three groups (Figure 4 panels B and D). Importantly, we found that the PEG-PAM treated animals had increased cortical capillary wall diameters compared to both the sham group (p=0.003) and the MCAO+Vehicle group (p=0.002). Capillary lumen diameter in the ipsilateral cortex was reduced compared to the contralateral cortex for both the MCAO+Vehicle (p<0.001) and MCAO+PAM (p<0.001) groups (Figure 4 panels C and E). While capillaries in the MCAO+PAM group had increased diameter compared to the MCAO+Vehicle group (p=0.006), this remained lower than the sham group (p=0.004).

**Figure 4.**
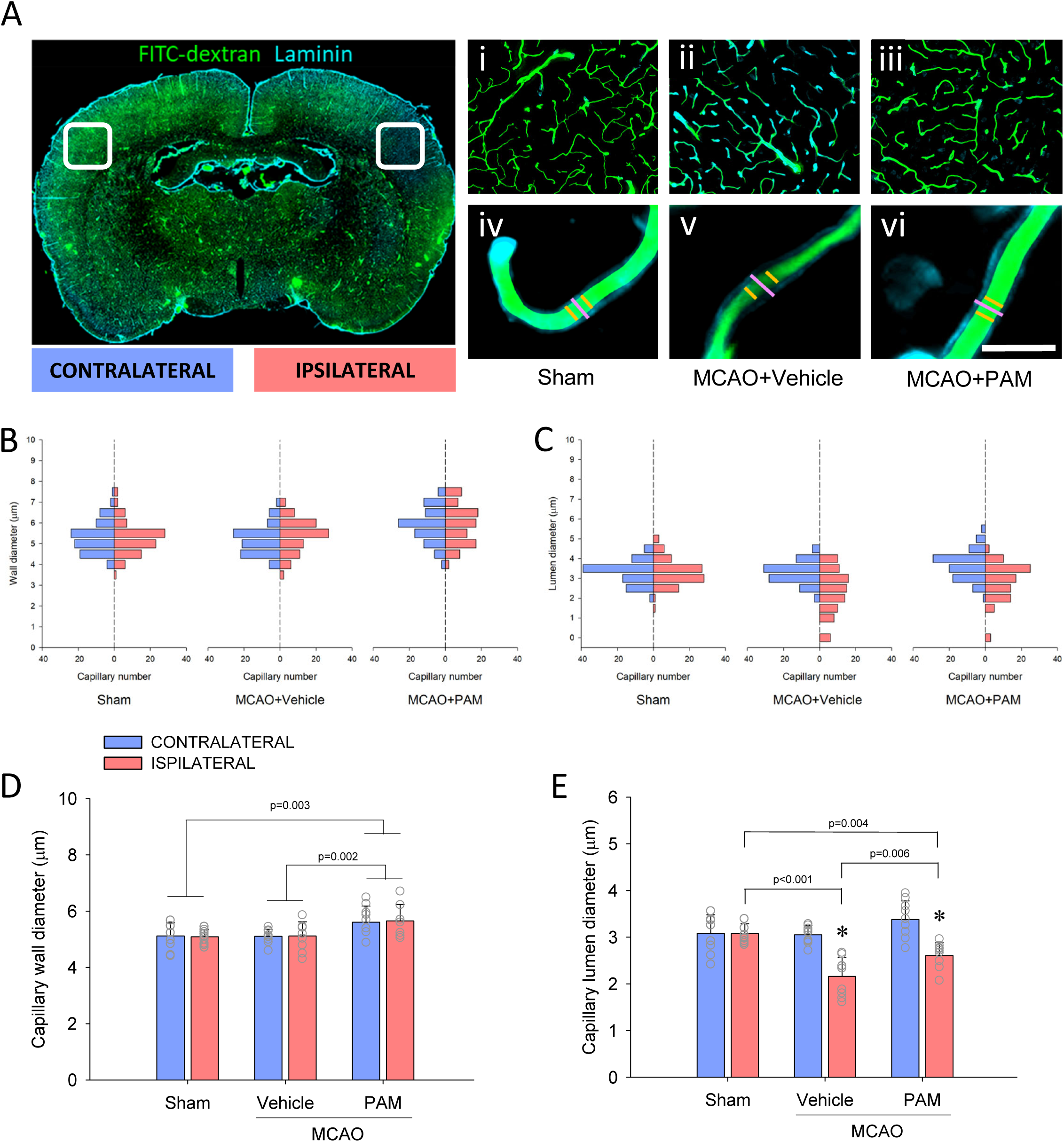
PAM stimulates capillary dilation and reduces capillary no-reflow in the cortex 24 hours after MCAO. Panel. **A.** Representative coronal brain section showing FITC-dextran (green) filling of vasculature (cyan) on the contralateral (no MCAO) and ipsilateral (MCAO) sides of the brain. White boxes indicate example regions of interest chosen for analysis of cortical capillaries. First row of insets (i-iii) show higher magnification examples of the cortical and striatal regions on both the contralateral and ipsilateral sides. Scale bar = 100µm. Second row of insets (iv-vi) show individual capillaries from the ipsilateral hemisphere with measurements of capillary wall diameter (pink line) and lumen diameter (orange line). Scale bar = 10µm. **Panels B and C.** Distribution frequencies for capillary wall diameter (B) and capillary lumen diameter (C) on the contralateral (blue) and ipsilateral (red) hemispheres. Measurements in each panel represent 10 individual capillaries from each animal in each group for both contralateral (n=80 capillaries) and ipsilateral (n=80 capillaries) hemispheres. **Panels D and E.** Quantification of mean capillary wall diameter (D) and capillary lumen diameter (E) from both contralateral and ipsilateral hemispheres. Data are means±SD for n=8 animals in each group. Open circles represent data from individual animals. Statistical analysis completed using two-way repeated measures ANOVA with Student-Newman Keuls post hoc. * denotes p<0.05 vs contralateral hemisphere within the group. Specific p values are provided for between group comparisons in all relevant panels.

**Figure 5.**
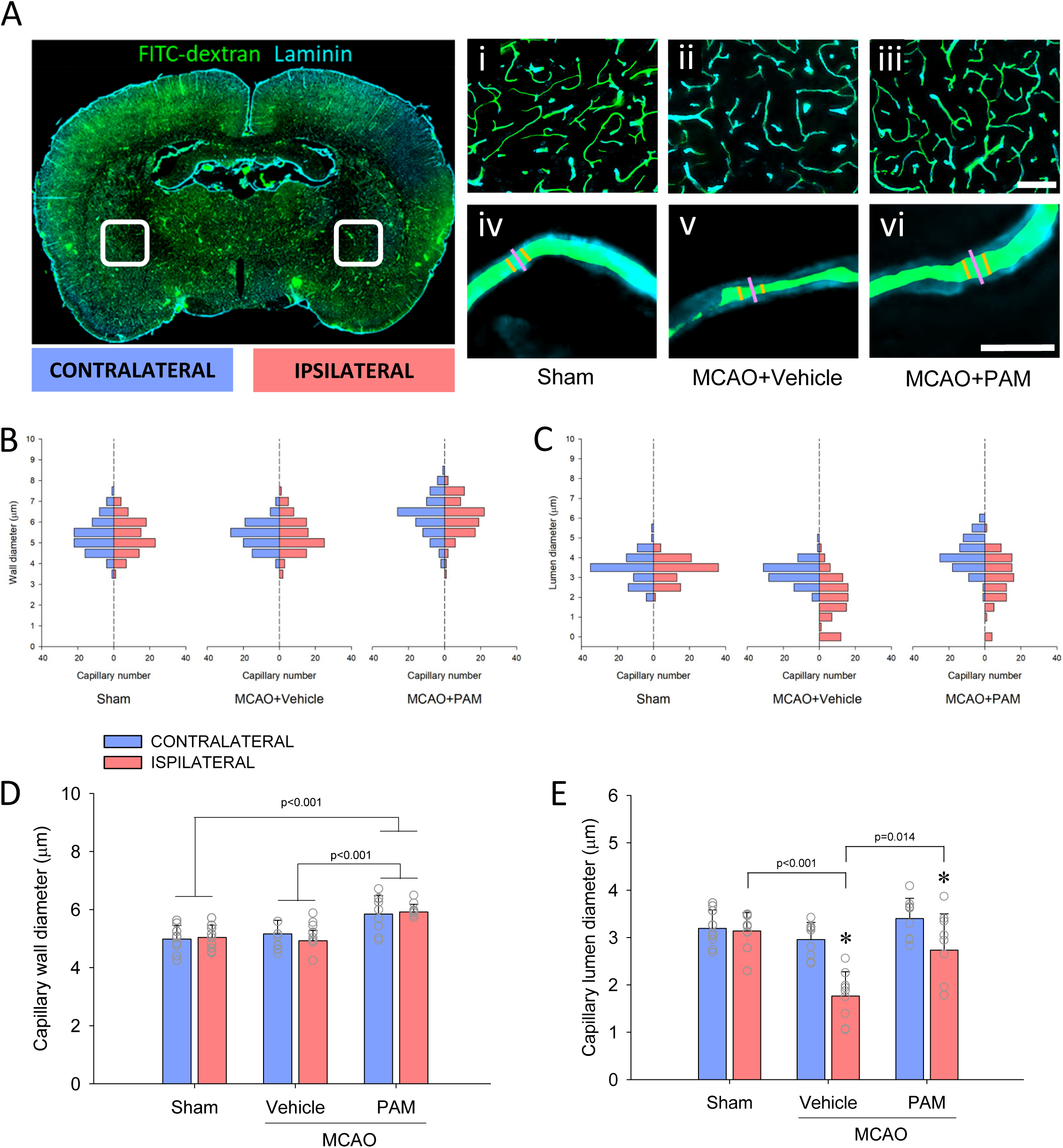
PAM stimulates capillary dilation and reduces capillary no-reflow in the subcortical region 24 hours after MCAO. Panel. **A.** Representative coronal brain section showing FITC-dextran (green) filling of vasculature (cyan) on the contralateral (no MCAO) and ipsilateral (MCAO) sides of the brain. White boxes indicate example regions of interest chosen for analysis of striatal capillaries. First row of insets (i-iii) show higher magnification examples of the cortical and striatal regions on both the contralateral and ipsilateral sides. Scale bar = 100µm. Second row of insets (iv-vi) show individual capillaries from the ipsilateral hemisphere with measurements of capillary wall diameter (pink line) and lumen diameter (orange line). Scale bar = 10µm. **Panels B and C.** Distribution frequency for capillary wall diameter (B) and capillary lumen diameter (C) on the contralateral (blue) and ipsilateral (red) hemispheres. Measurements in each panel represent 10 individual capillaries from each animal in each group for both contralateral (n=80 capillaries) and ipsilateral (n=80 capillaries) hemispheres. **Panels D and E.** Quantification of mean capillary wall diameter (D) and capillary lumen diameter (E) from both contralateral and ipsilateral hemispheres. Data are means±SD for n=8 animals in each group. Open circles represent data from individual animals. Statistical analysis was completed using two-way repeated measures ANOVA with Student-Newman Keuls post hoc. * denotes p<0.05 vs contralateral hemisphere within the group. Specific p values are provided for between group comparisons in all relevant panels.

Next, we assessed whether changes in capillary wall and lumen diameters in the striatal region followed a similar trend to that seen in the cortex (Figure 5). Capillary wall diameters within the striatum did not differ between the contralateral and ipsilateral hemispheres for all three groups (Figure 5 panels B and D). As in the cortex, we found that striatal capillary wall diameters within the MCAO+PAM group were significantly larger than the sham group (p<0.001) and the MCAO+Vehicle group (p<0.001). Striatal capillary lumen diameter within the ipsilateral hemisphere was much smaller than in the contralateral hemisphere for both the vehicle-treated (p<0.001) and PEG-PAM-treated (p=0.005) groups (Figure 5 panels C and E). Capillary lumen diameter within the ipsilateral striatum was increased in the MCAO+PAM group compared to the MCAO+Vehicle group (p=0.014). Importantly, we found that ipsilateral striatal capillary lumen diameters for the MCAO+PAM group were similar to that of the sham group (p=0.080).

### PAM improves brain blood flow and reduces neurological deficit and infarct volume 7 days after MCAO

In the second experiment, investigated whether the acute improvements in blood flow after PEG-PAM administration led to improved longer-term recovery of the brain following MCAO. Changes in vascular perfusion before, during and immediately after MCAO within cortical and subcortical regions were quantified using tCEU, as in experiment 1. As expected, the sham group showed no significant differences in cortical or subcortical vascular perfusion across any time-point (Figure 6 panels A and B). Vascular perfusion within the cortex was found to decrease upon the induction of MCAO for both the vehicle-treated and PAM-treated groups (Figure 6A; p<0.001 for both). Cortical vascular perfusion then returned to baseline following reperfusion for both the MCAO+Vehicle and MCAO+PAM groups. In line with experiment 1, no differences in cortical vascular perfusion were found between the MCAO+Vehicle and MCAO+PAM groups across any of the three time-points.

**Figure 6.**
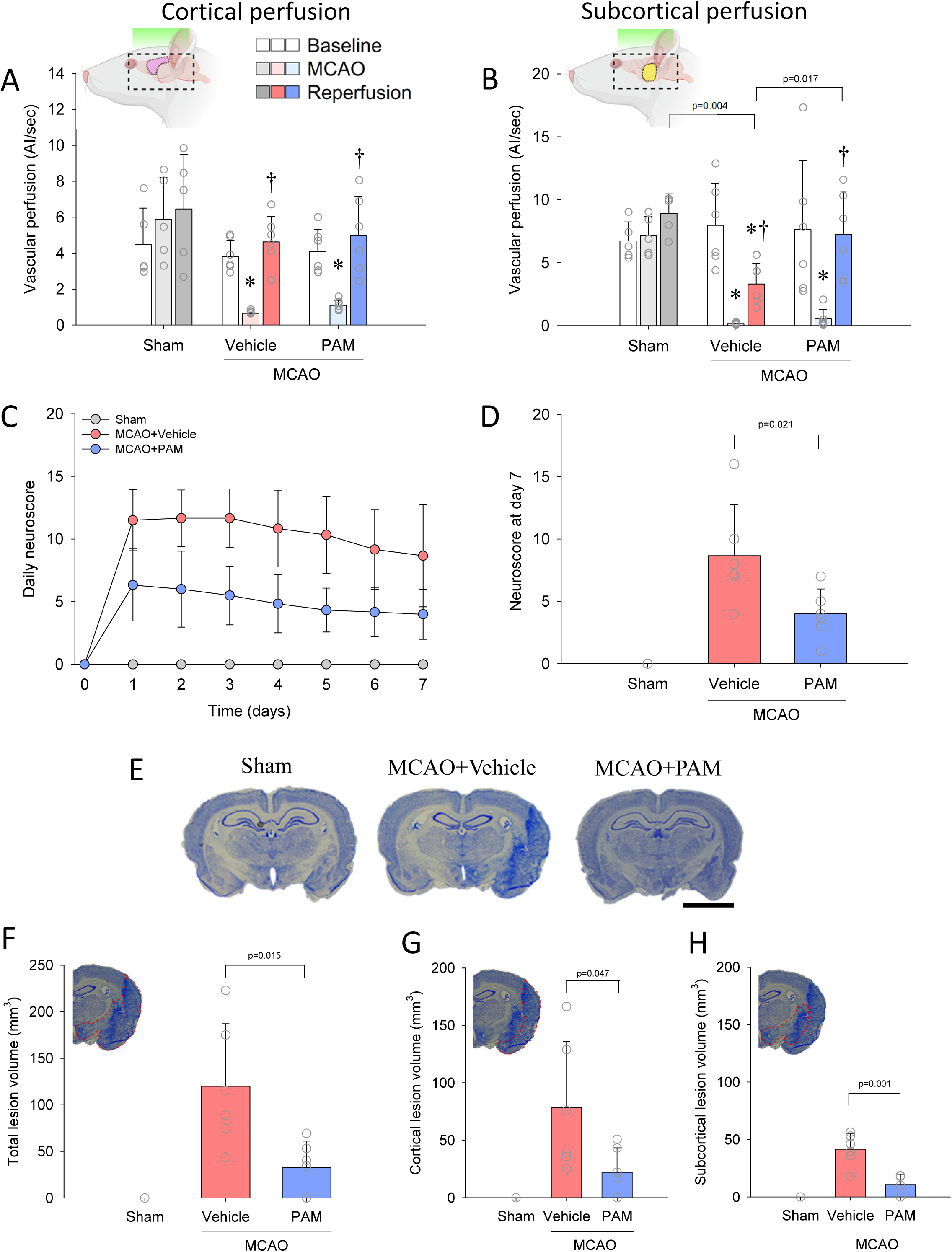
PAM improves brain blood flow at acute reperfusion and reduces neurological deficit and infarct volume 7 days after MCAO. Panel. **A.** Quantification of cortical vascular perfusion at baseline, MCAO, and acute reperfusion across all groups. **Panel B.** Quantification of subcortical vascular perfusion at baseline, MCAO, and acute reperfusion across all groups. **Panels C and D**. Daily neuroscore assessment for each group across the 7-day recovery period (C) with quantified data for day 7 shown in panel D. **Panel E.** Representative images of coronal brain sections stained with Cresyl Violet from each group. Scale bar = 5mm. **Panels F-H.** Quantification of total (F), cortical (G) and subcortical (H) infarct volumes. Data are means±SD for Sham (n=5), MCAO+Vehicle (n=6) and MCAO+PAM (n=6) groups. Open circles represent data from individual animals. Statistical analysis for panels A-C was completed using two-way repeated measures ANOVA with Student-Newman Keuls post hoc. One-way ANOVA with Student-Newman Keuls post hoc was used for analysis of data in panels D-G. The Kruskal-Wallis test was used to analyze neuroscore data in panel D. * denotes p<0.05 vs baseline within the group. † denotes p<0.05 vs MCAO within the group. Specific p values are provided for between group comparisons in all relevant panels.

The induction of MCAO caused a significant drop in subcortical vascular perfusion for both the vehicle-treated and PEG-PAM-treated groups (Figure 6B; p<0.001 for both). However, upon reperfusion, subcortical vascular perfusion remained significantly lower than baseline for the MCAO+Vehicle group (p<0.001). Conversely, subcortical vascular perfusion within the MCAO+PAM group returned to baseline following reperfusion mirroring the effects observed in experiment 1. At reperfusion, the MCAO+PAM group had higher vascular perfusion within the subcortex compared to the vehicle-treated group (p=0.017).

Next, we assessed whether the improvement in acute blood flow after PEG-PAM administration impacted longer-term neurological recovery following MCAO. To do this, we performed daily neuroscore analyses for 7 days following MCAO. As expected, animals in the sham group showed no neurological deficit over the 7-day recovery period (Figure 6C). In contrast, animals in the MCAO+Vehicle group exhibited marked neurological impairment compared to sham. Importantly, we found that animals in the MCAO+PAM group exhibited ∼50% lower score for neurological deficit compared to the MCAO+Vehicle group across all 7 days (p<0.001 for treatment effect). The day 7 neuroscore for the PEG-PAM-treated group was approximately 55% lower than that of the vehicle-treated group (Figure 6D; p=0.021).

We next determined if the observed improvement in neurological recovery coincides with a reduction in the extent of brain damage caused by ischemia. To do this, infarct volume was quantified using cresyl violet staining (Figure 6E), as in experiment 1. As expected, the sham group had no identifiable infarcts in the brain (Figure 6F). After the 7-day recovery period, we found that the MCAO+PAM group had markedly reduced total lesion volume than the MCAO+Vehicle group (p=0.015). In line with this and our vascular perfusion data, we found that the PAM treatment after MCAO reduced infarct volume in both the cortex (Figure 6G; p=0.047) and subcortex (Figure 6H; p=0.001) when compared to the vehicle treatment.

### PEG-PAM administration leads to elevated circulating PEG-PAM 24 hours after ischemic stroke

Lastly, we quantified circulating PEG-PAM levels at the 24-hour endpoint to confirm target engagement. The sham (0.03 ± 0.18 ng/ml) and MCAO+Vehicle (0.06 ± 0.17 ng/ml) groups had no detectable circulating PEG-PAM, while animals treated with PEG-PAM showed markedly elevated circulating PEG-PAM concentrations 24 hours after a single intraperitoneal administration (255 ± 124 ng/ml; p<0.001 vs both other groups).

## DISCUSSION

The primary goal of this study was to determine whether PEG-PAM administration could improve microvascular function, reduce capillary no-reflow and enhance brain recovery after ischemic stroke. We show that PEG-PAM administration in the acute phase after ischemic stroke leads to improved microvascular blood flow for at least the following 24 hours and that this is associated with enhanced neurological recovery across the following 7 days. The persistence of these effects beyond the acute ischemic phase supports sustained PAM-mediated peptide amidation as a potential therapeutic strategy to augment brain recovery after ischemic stroke.

Microvascular dysfunction and development of capillary no-reflow is common and proposed to contribute to ongoing brain ischemia and damage following ischemic stroke, even after intervention to restore arterial blood flow (9). While multiple mechanisms drive microvascular defects after ischemia, capillary no-reflow has been identified as a major contributor to ongoing brain damage and increased risk of death after ischemic stroke (3, 9, 10). We used *in vivo* imaging and post-mortem analysis to show that capillary no-flow impacts large regions of the rodent brain for at least 24 hours after ischemic stroke. These results are in line with previous studies highlighting capillary no-reflow as a common feature of the damaged brain tissue after ischemia (5, 43, 44). Importantly, we also show that PEG-PAM administration leads to capillary dilation and restoration of blood flow following ischemia and that this is associated with improved neurological recovery over the following 7 days. In this context, our findings suggest that enhancing microvascular blood flow after ischemic stroke represents a critical therapeutic target beyond restoration of large vessel patency. This outcome is particularly relevant given that microvascular blood flow and its dysfunction after ischemic stroke remain largely unaddressed by current reperfusion therapies which only target arterial recanalization.

The precise mechanism by which PEG-PAM improves capillary perfusion and neurological recovery remains to be determined. PAM-mediated α-amidation is required for the biological activity of numerous vasoactive and neuroprotective peptides (12, 13), and the beneficial effects observed in the present study are likely mediated through augmentation of several of these endogenous pathways (27). Importantly, our findings support the concept that therapeutic augmentation of circulating PAM activity can modify vascular recovery and improve capillary perfusion following ischemic injury in the acute stage (24 hours) that subsequently improves brain recovery in the longer term. Together with previous studies showing that recombinant PAM remains catalytically active in blood and converts glycine-extended peptide precursors into their amidated products *in vivo* (27), the present findings further support the circulation as a therapeutically accessible compartment for peptide amidation.

It is important to note that PAM-dependent peptide systems are differentially regulated following ischemic stroke. While some amidated peptides with vasoprotective and neuroprotective functions are decreased following cerebral ischemia (21, 45), others, including ADM, copeptin/vasopressin and Substance P, are elevated (14, 46, 47) and have been proposed to be biomarkers of stroke severity where increased vascular damage drives increased secretion of these vasoprotective hormones as a compensatory response. For this reason, others have investigated whether acutely increasing bioactive ADM after stroke via intravenous infusion can improve brain recovery (19, 23). While this approach did not markedly modify recovery after stroke, potentialy due to limited half life of ADM, the infusion of ADM was well tolerated with no adverse effects noted. Similarly, in our study PEG-PAM administration did not induce systemic hypotension or other adverse hemodynamic effects in healthy rats, supporting the safety and translational potential of this approach. Unlike administration of a single vasoactive peptide at supraphysiological concentrations (23), PEG-PAM enhances the conversion of endogenous glycine-extended precursors into their bioactive forms and therefore preserves the physiological regulation of peptide release and signalling. Therefore, this approach may be more relevant to improving vascular recovery following injury, such as ischemia. In support of this, recent studies indicate that the balance between amidated peptides and their glycine-extended precursors may be more informative than absolute peptide concentrations alone (48). Thiele et al. (2025) demonstrated that the ratio between bio-ADM and ADM-Gly reflects pathway activation more comprehensively than either analyte individually (49). Furthermore, experimental studies in porcine sepsis showed that increased PAM enhances conversion of ADM-Glyine into bioactive ADM, providing direct evidence that peptide amidation can occur within the circulation (49). These observations suggest that PAM augmentation may influence vascular biology by improving the efficiency of endogenous peptide maturation and shifting the balance between inactive precursor pools and biologically active amidated products, rather than simply increasing peptide abundance. Importantly, sustained PAM activity over the 7-day post-PEG-PAM administration window is consistent with continued amidating capacity well beyond the acute (24 hour) ischemic phase. While the precise molecular pathways activated were not directly investigated in our study, the durable improvement in capillary perfusion, infarct and behavioral recovery may be explained by acute-phase PAM-mediated peptide activation establishing microvascular reperfusion, with downstream consequences extending across the 7-day recovery window. The specific peptites activated by PAM that are important for vascular and brain recovery after ischemic stroke warrant future investigation.

There are several limitations associated with this study. Firstly, we did not measure the temporal profile of PAM-mediated peptide amidation following PEG-PAM administration in the post-stroke setting. Longitudinal blood sampling at the volumes required would have substantially impacted recovery of the animals after induction of ischemic stroke. Given we have previously shown that PEG-PAM remains in the circulation for up to 7 days after administration (28), it is reasonable to conclude that PAM activity remains elevated beyond the acute phase of the present study and contributes to microvascular and brain recovery over the following 7 days. Secondly, although we have assessed microvascular blood flow and capillary diameters directly, we have not identified the specific PAM substrate(s) responsible for the observed vascular benefit. Bioactive ADM has been reported to exert protective effects in ischemia through nitric-oxide-mediated vasodilation, improved barrier function, reduced edema, reduced oxidative stress and reduced inflammation (45, 46). However, ADM is unlikely to be the sole mediator of the effects observed in the present study. Other PAM-dependent peptides, including CGRP, VIP and PACAP, possess similar vasodilatory, anti-inflammatory and neuroprotective properties and have demonstrated beneficial effects in experimental stroke models (50, 51). Consequently, augmentation of PAM activity may provide a broader and potentially more physiologically relevant response than administration of any individual peptide alone. Further work is required to resolve which of these mechanisms predominantly drives the improvement in vascular and brain recovery after ischemia.

In conclusion, the results of this study highlight that PEG-PAM administration improves microvascular function and reduces capillary no-reflow to aid brain recovery after ischemic stroke. This is one of the first studies highlighting that improvement in microvascular function and reduction of capillary no-reflow acutely after ischemia can enhance longer-term brain recovery. Therefore, our work may represent a novel therapeutic avenue for reducing brain damage and improving functional recovery after ischemic stroke.

## Acknowledgments

None.

## Author Contributions

DP, YI and PK contributed to the conceptualization of the study. DP, EM, YI, AT, EA, SM contributed to the methodology and data collection. All authors contributed to data analysis, visualization and interpretation. EM, YI and DP drafted the manuscript and all authors contributed to reviewing and editing of the manuscript.

## REFERENCES

1. Goyal M, Menon BK, Van Zwam WH, Dippel DW, Mitchell PJ, Demchuk AM, et al. Endovascular thrombectomy after large-vessel ischaemic stroke: a meta-analysis of individual patient data from five randomised trials. The Lancet. 2016;387(10029):1723–31.

2. Sarraj A, Hassan AE, Abraham MG, Ortega-Gutierrez S, Kasner SE, Hussain MS, et al. Trial of endovascular thrombectomy for large ischemic strokes. New England Journal of Medicine. 2023;388(14):1259–71.

3. Ng FC, Churilov L, Yassi N, Kleinig TJ, Thijs V, Wu TY, et al. Microvascular dysfunction in blood-brain barrier disruption and hypoperfusion within the infarct posttreatment are associated with cerebral edema. Stroke. 2022;53(5):1597–605.

4. Tudor T, Spinazzi EF, Alexander JE, Mandigo GK, Lavine SD, Grinband J, et al. Progressive microvascular failure in acute ischemic stroke: A systematic review, meta-analysis, and time-course analysis. Journal of Cerebral Blood Flow & Metabolism. 2024;44(2):192–208.

5. Hall CN, Reynell C, Gesslein B, Hamilton NB, Mishra A, Sutherland BA, et al. Capillary pericytes regulate cerebral blood flow in health and disease. Nature. 2014;508(7494):55–60.

6. El Amki M, Glueck C, Binder N, Middleham W, Wyss MT, Weiss T, et al. Neutrophils obstructing brain capillaries are a major cause of no-reflow in ischemic stroke. Cell reports. 2020;33(2).

7. Cancino A, Muñoz P, Cox P, Acevedo L, Castillo S, Letelier A, et al. Effect of inflammation on neurovascular coupling, microperfusion, and clinical outcomes in ischemic stroke patients: a case series report. Frontiers in Medicine. 2025;12:1665396.

8. Hu J, Nan D, Lu Y, Niu Z, Ren Y, Qu X, et al. Microcirculation no-reflow phenomenon after acute ischemic stroke. European neurology. 2023;86(2):85–94.

9. Ng FC, Churilov L, Yassi N, Kleinig TJ, Thijs V, Wu T, et al. Prevalence and significance of impaired microvascular tissue reperfusion despite macrovascular angiographic reperfusion (no-reflow). Neurology. 2022;98(8):e790–e801.

10. Hussein HM, Saleem MA, Qureshi AI. Rates and predictors of futile recanalization in patients undergoing endovascular treatment in a multicenter clinical trial. Neuroradiology. 2018;60:557–63.

11. Pereira da Silva AM, Ribeiro Gonçalves O, Falcão L, Ribeiro FV, Han ML, Rodrigues Menezes I, et al. Association between the no-reflow phenomenon and clinical outcomes after endovascular treatment for acute ischemic stroke: A systematic review and meta-analysis. European Stroke Journal. 2025:23969873251376846.

12. Eipper BA, Stoffers DA, Mains RE. The biosynthesis of neuropeptides: peptide alpha-amidation. Annu Rev Neurosci. 1992;15:57–85.

13. Eipper BA, Milgram SL, Jean Husten E, Yun HY, Mains RE. Peptidylglycine α-amidating monooxygenase: A multifunctional protein with catalytic, processing, and routing domains. Protein Science. 1993;2(4):489–97.

14. Zhang H, Tang B, Yin C-G, Chen Y, Meng Q-L, Jiang L, et al. Plasma adrenomedullin levels are associated with long-term outcomes of acute ischemic stroke. Peptides. 2014;52:44–8.

15. Bahr-Hosseini M, Meissner N, Reidler P, Saver JL, Tiedt S. Plasma CGRP Levels Are Not Associated With Collateral Flow and Outcome After Stroke. Stroke. 2023;54(5):e203–e4.

16. Hippenstiel S, Witzenrath M, Schmeck B, Hocke A, Krisp M, Krüll M, et al. Adrenomedullin reduces endothelial hyperpermeability. Circulation research. 2002;91(7):618–25.

17. Kis B, Kaiya H, Nishi R, Deli M, Abraham C, Yanagita T, et al. Cerebral endothelial cells are a major source of adrenomedullin. Journal of neuroendocrinology. 2002;14(4):283–93.

18. Uddman R, Goadsby PJ, Jansen I, Edvinsson L. PACAP, a VIP-like peptide: immunohistochemical localization and effect upon cat pial arteries and cerebral blood flow. J Cereb Blood Flow Metab. 1993;13(2):291–7.

19. Xia C-F, Yin H, Borlongan CV, Chao J, Chao L. Postischemic infusion of adrenomedullin protects against ischemic stroke by inhibiting apoptosis and promoting angiogenesis. Experimental neurology. 2006;197(2):521–30.

20. Chaung WW, Wu R, Ji Y, Wang Z, Dong W, Cheyuo C, et al. Peripheral administration of human adrenomedullin and its binding protein attenuates stroke-induced apoptosis and brain injury in rats. Molecular Medicine. 2011;17(9-10):1075–83.

21. Chen G, Frokiaer J, Pedersen M, Nielsen S, Si Z, Pang Q, et al. Reduction of ischemic stroke in rat brain by alpha melanocyte stimulating hormone. Neuropeptides. 2008;42(3):331–8.

22. Meeran K, O’Shea D, Upton PD, Small CJ, Ghatei MA, Byfield PH, et al. Circulating adrenomedullin does not regulate systemic blood pressure but increases plasma prolactin after intravenous infusion in humans: a pharmacokinetic study. The Journal of Clinical Endocrinology & Metabolism. 1997;82(1):95–100.

23. Yoshimoto T, Saito S, Omae K, Tanaka K, Kita T, Kitamura K, et al. Efficacy and safety of adrenomedullin for acute ischemic stroke (AMFIS): a phase 2, randomized, double-blinded, placebo-controlled, clinical trial. EClinicalMedicine. 2024;77.

24. Eipper BA, Mains RE, Glembotski CC. Identification in pituitary tissue of a peptide alpha-amidation activity that acts on glycine-extended peptides and requires molecular oxygen, copper, and ascorbic acid. Proc Natl Acad Sci U S A. 1983;80(16):5144–8.

25. Kaufmann P, Ilina Y, Press M, Bergmann A. Sandwich immunoassay for adrenomedullin precursor and its practical application. Sci Rep. 2024;14(1):28091.

26. Siddheshwar RK, Gray JC, Kelly SB. Plasma levels of progastrin but not amidated gastrin or glycine extended gastrin are elevated in patients with colorectal carcinoma. Gut. 2001;48(1):47–52.

27. Kaufmann P, Bergmann A, Melander O. Novel insights into peptide amidation and amidating activity in the human circulation. Scientific Reports. 2021;11(1):15791.

28. Ilina Y, Kaufmann P, Press M, Uba TI, Bergmann A. Enhancing stability and bioavailability of peptidylglycine alpha-amidating monooxygenase in circulation for clinical use. Biomolecules. 2025;15(2):224.

29. Ilina Y, Kaufmann P, Melander O, Press M, Thuene K, Bergmann A. Immunoassay-based quantification of full-length peptidylglycine alpha-amidating monooxygenase in human plasma. Sci Rep. 2023;13(1):10827.

30. Premilovac D, Blackwood SJ, Ramsay CJ, Keske MA, Howells DW, Sutherland BA. Transcranial contrast-enhanced ultrasound in the rat brain reveals substantial hyperperfusion acutely post-stroke. Journal of Cerebral Blood Flow & Metabolism. 2020;40(5):939–53.

31. Sutherland BA, Neuhaus AA, Couch Y, Balami JS, DeLuca GC, Hadley G, et al. The transient intraluminal filament middle cerebral artery occlusion model as a model of endovascular thrombectomy in stroke. Journal of Cerebral Blood Flow & Metabolism. 2016;36(2):363–9.

32. Paxinos G, Watson C. The rat brain in stereotaxic coordinates: hard cover edition: Elsevier; 2006.

33. Ng HL, Premilovac D, Rattigan S, Richards SM, Muniyappa R, Quon MJ, et al. Acute vascular and metabolic actions of the green tea polyphenol epigallocatechin 3-gallate in rat skeletal muscle. The Journal of Nutritional Biochemistry. 2017;40:23–31.

34. Russell RD, Roberts-Thomson KM, Hu D, Greenaway T, Betik AC, Parker L, et al. Impaired postprandial skeletal muscle vascular responses to a mixed meal challenge in normoglycaemic people with a parent with type 2 diabetes. Diabetologia. 2022;65(1):216–25.

35. Hu D, Remash D, Russell RD, Greenaway T, Rattigan S, Squibb KA, et al. Impairments in adipose tissue microcirculation in type 2 diabetes mellitus assessed by real-time contrast-enhanced ultrasound. Circulation: Cardiovascular Imaging. 2018;11(4):e007074.

36. Encarnacion A, Horie N, Keren-Gill H, Bliss TM, Steinberg GK, Shamloo M. Long-term behavioral assessment of function in an experimental model for ischemic stroke. Journal of neuroscience methods. 2011;196(2):247–57.

37. Garcia JH, Wagner S, Liu K-F, Hu X-j. Neurological deficit and extent of neuronal necrosis attributable to middle cerebral artery occlusion in rats: statistical validation. Stroke. 1995;26(4):627–35.

38. Saver JL, Chaisinanunkul N, Campbell BC, Grotta JC, Hill MD, Khatri P, et al. Standardized nomenclature for modified Rankin scale global disability outcomes: consensus recommendations from stroke therapy academic industry roundtable XI. Stroke. 2021;52(9):3054–62.

39. Attrill E, Richards SM, Ross RM, Sutherland BA, Premilovac D. Induction of type 2 diabetes in mice to understand vascular changes that drive diabetic retinopathy. Diabetic Retinopathy: Methods and Protocols: Springer; 2023. p. 1–12.

40. Bankhead P, Loughrey MB, Fernández JA, Dombrowski Y, McArt DG, Dunne PD, et al. QuPath: Open source software for digital pathology image analysis. Scientific reports. 2017;7(1):1–7.

41. Nouraee C, Fisher M, Di Napoli M, Salazar P, Farr TD, Jafarli A, et al. A brief review of edema-adjusted infarct volume measurement techniques for rodent focal cerebral ischemia models with practical recommendations. Journal of vascular and interventional neurology. 2019;10(3):38.

42. Box GEP, Cox DR. An Analysis of Transformations. Journal of the Royal Statistical Society: Series B (Methodological). 1964;26(2):211–43.

43. Beard DJ, Brown LS, Morris GP, Couch Y, Adriaanse BA, Karali CS, et al. Rapamycin Treatment Reduces Brain Pericyte Constriction in Ischemic Stroke. Translational Stroke Research. 2025;16(4):1185–97.

44. Sun F, Zhou J, Chen X, Yang T, Wang G, Ge J, et al. No-reflow after recanalization in ischemic stroke: from pathomechanisms to therapeutic strategies. Journal of Cerebral Blood Flow & Metabolism. 2024;44(6):857–80.

45. Zierath D, Tanzi P, Cain K, Shibata D, Becker K. Plasma alpha-melanocyte stimulating hormone predicts outcome in ischemic stroke. Stroke. 2011;42(12):3415–20.

46. Katan M, Fluri F, Morgenthaler NG, Schuetz P, Zweifel C, Bingisser R, et al. Copeptin: a novel, independent prognostic marker in patients with ischemic stroke. Ann Neurol. 2009;66(6):799–808.

47. Lorente L, Martin MM, Almeida T, Perez-Cejas A, Ramos L, Argueso M, et al. Serum Levels of Substance P and Mortality in Patients with a Severe Acute Ischemic Stroke. Int J Mol Sci. 2016;17(6).

48. Marino R, Struck J, Maisel AS, Magrini L, Bergmann A, Di Somma S. Plasma adrenomedullin is associated with short-term mortality and vasopressor requirement in patients admitted with sepsis. Crit Care. 2014;18(1):R34.

49. Thiele C, Ilina Y, Kaufmann P, Springsfeld G, Press M, Hartmann O, et al. Kinetics of adrenomedullin pathway activation in a porcine sepsis model and a human cohort of sepsis and septic shock. Scientific Reports. 2025;15(1):32693.

50. Reglodi D, Somogyvari-Vigh A, Vigh S, Kozicz T, Arimura A. Delayed systemic administration of PACAP38 is neuroprotective in transient middle cerebral artery occlusion in the rat. Stroke. 2000;31(6):1411–7.

51. Yang J, Zong CH, Zhao ZH, Hu XD, Shi QD, Xiao XL, et al. Vasoactive intestinal peptide in rats with focal cerebral ischemia enhances angiogenesis. Neuroscience. 2009;161(2):413–21.

