## Supplementary material for "Peptidylglycine α-amidating monooxygenase restores brain microvascular blood flow and improves recovery following ischemic stroke"

* denotes equal first authors

Key words: ischemic stroke, capillary no reflow, peptidylglycine α-amidating monooxygenase, contrast enhanced ultrasound, brain blood flow, amidated peptides

Running title: PAM reduces capillary no reflow 24 hours after ischemic stroke.

Corresponding author:

Dino Premilovac (PhD)

Tasmanian School of Medicine

Health, University of Tasmania

Hobart, TAS 7000, Australia

Ph: +61 3 6226 2701

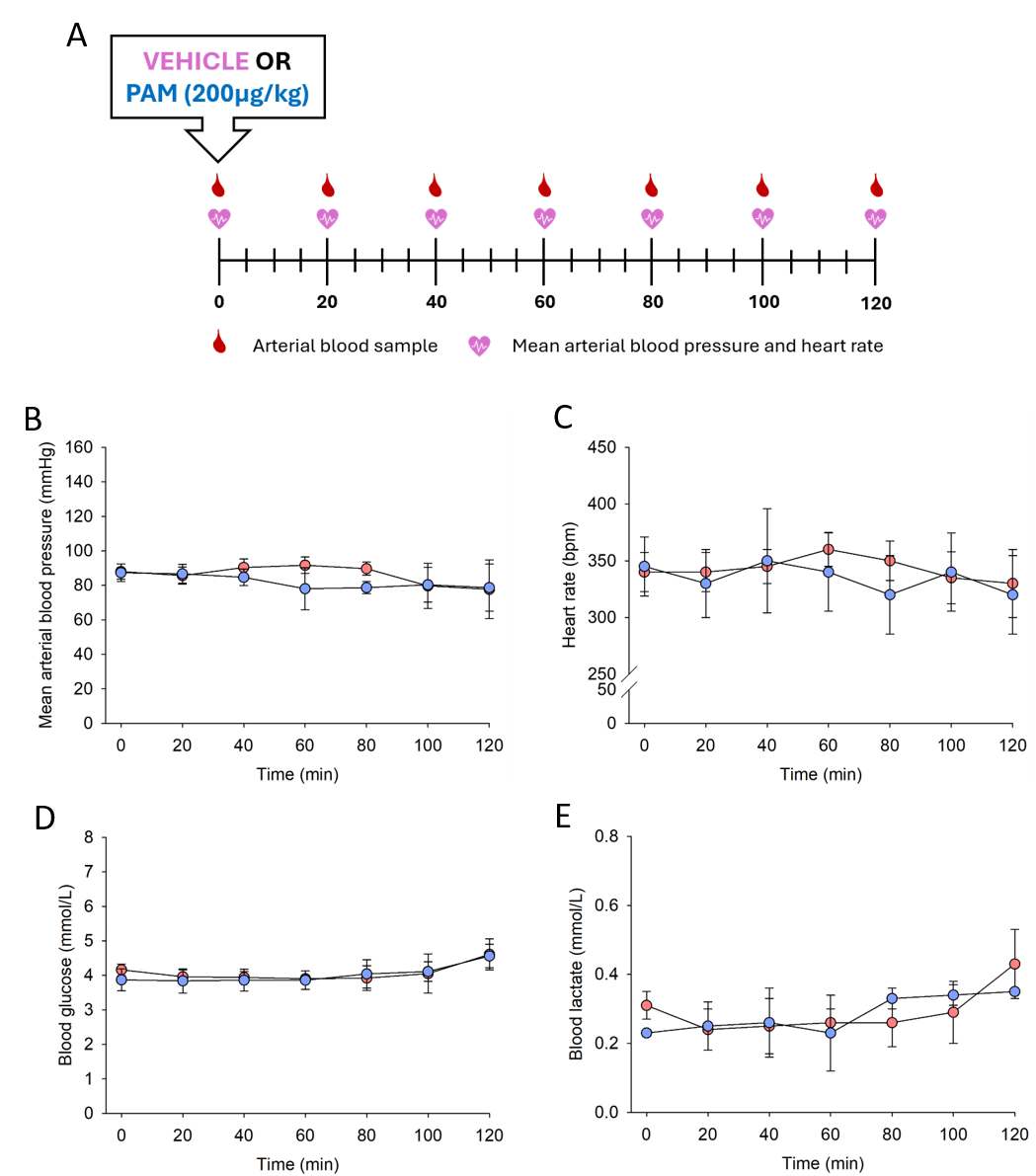

**Supplementary Figure 1. PAM administration does not alter systemic hemodynamics or metabolism in healthy rats. A.** Visual representation of the time course for the experiment. All rats were anesthetized with pentobarbitone (85mg/kg) prior to undergoing minor surgery to cannulate the carotid artery and both jugular veins. At Time = 0 minutes, rats received an intraperitoneal injection of either a PAM (200µg/kg) or vehicle (saline). Arterial blood samples and heart rate and blood pressure recordings were collected every 20 minutes from Time = 0 minutes. After 120 minutes, rats were euthanized with an intravenous overdose of pentobarbitone (>200mg/kg). **Panels B-E.** Time course measurements of mean arterial blood pressure (B), heart rate (C), blood glucose (D) and blood lactate (E) after PAM administration. Data are means±SD for n=3 animals in each group.
